# High-throughput Multimodal Automated Phenotyping (MAP) with Application to PheWAS

**DOI:** 10.1101/587436

**Authors:** Katherine P. Liao, Jiehuan Sun, Tianrun A. Cai, Nicholas Link, Chuan Hong, Jie Huang, Jennifer E. Huffman, Jessica Gronsbell, Yichi Zhang, Yuk-Lam Ho, Victor Castro, Vivian Gainer, Shawn N. Murphy, Christopher J. O’Donnell, J. Michael Gaziano, Kelly Cho, Peter Szolovits, Isaac S. Kohane, Sheng Yu, Tianxi Cai, with the VA Million Veteran Program

## Abstract

**Objective:** Electronic health records (EHR) linked with biorepositories are a powerful platform for translational studies. A major bottleneck exists in the ability to phenotype patients accurately and efficiently. The objective of this study was to develop an automated high-throughput phenotyping method integrating International Classification of Diseases (ICD) codes and narrative data extracted using natural language processing (NLP).

**Method:** We developed a mapping method for automatically identifying relevant ICD and NLP concepts for a specific phenotype leveraging the UMLS. Aggregated ICD and NLP counts along with healthcare utilization were jointly analyzed by fitting an ensemble of latent mixture models. The MAP algorithm yields a predicted probability of phenotype for each patient and a threshold for classifying subjects with phenotype yes/no. The algorithm was validated using labeled data for 16 phenotypes from a biorepository and further tested in an independent cohort PheWAS for two SNPs with known associations.

**Results:** The MAP algorithm achieved higher or similar AUC and F-scores compared to the ICD code across all 16 phenotypes. The features assembled via the automated approach had comparable accuracy to those assembled via manual curation (AUC_MAP_ 0.943, AUC_manual_ 0.941). The PheWAS results suggest that the MAP approach detected previously validated associations with higher power when compared to the standard PheWAS method based on ICD codes.

**Conclusion:** The MAP approach increased the accuracy of phenotype definition while maintaining scalability, facilitating use in studies requiring large scale phenotyping, such as PheWAS.

## INTRODUCTION

High-throughput technologies have provided powerful tools for dissecting the genomic and biologic architecture of complex human traits. To continue on the trajectory of their success, more work is needed to translate ‘omics’ findings to improvement in patient care. A major challenge in realizing such translation is linking the high-throughput biologic data with detailed high-quality phenotypic data. To improve their translational potential, ‘omics’ studies are increasingly conducted using cohorts with linked Electronic Health Records (EHR) data and biobanks. Such studies are typically designed to investigate hundreds to thousands of phenotypes simultaneously, identifying a large unmet need for scalable, standardized, efficient and portable approaches for phenotyping.

The current standard for studying hundreds to thousands of phenotypes over millions of patients is using diagnoses codes such as the International Classification of Diseases (ICD) codes. The Phenome-Wide Association Studies (PheWAS) approach[1] was developed specifically for use in EHRs linked with biobanks to screen for associations between a genetic marker with roughly 1800 phenotypes. The phenotypes in the PheWAS are generated by grouping ICD9 codes. However, a known pitfall of ICD codes are variations in accuracy, leading to loss of power for association studies[2-5]. Rule-based manual curation and machine learning-based supervised learning approaches have been proposed as more accurate alternatives for phenotyping and have been used to create EHR-based patient cohorts for discovery research[6-18]. These approaches enhance the accuracy of phenotyping by combining information from multiple EHR features, including codified features such as ICD codes and narrative features identified via natural language processing (NLP). While collaborative platforms such as the PheKB[19] share existing phenotyping algorithms, deploying these algorithms across institutions can be labor intensive, limiting the feasibility of using these algorithms for phenotype screens.

To improve the efficiency for developing phenotyping algorithms, various approaches have been developed to reduce the level of human input required. Examples of these approaches include active sampling, feature refinement, and feature selection[20-23]. As well, several automated annotation methods have also been proposed[24 25]. However, these methods either still require expert input to curate silver standard labels or have variable accuracy. Recently, Yu et al., developed ‘PheNorm’[26], a fully automated phenotyping algorithm based on normal mixture modeling using ICD codes only. While PheNorm presented a highly scalable approach by ranking subjects with respect to their likelihood of having a phenotype, it could not provide the final classification of whether a subject had a specific phenotype necessary for clinical studies.

In this paper, we propose an unsupervised multimodal automated phenotyping (MAP) method with application in PheWAS. We hypothesize that MAP, which incorporates NLP, will efficiently and accurately classify millions of subjects across the roughly 1800 phenotypes in the PheWAS compared to existing approaches.

## METHODS

The MAP procedure includes two key steps: (i) assembling the ICD codes and NLP features corresponding to the target phenotype; (ii) annotation via unsupervised ensemble latent mixture modeling.

### Identifying the Main ICD and NLP Features for Phenotypes

For a target phenotype, we first identify the main ICD code associated with the phenotype. The main ICD code is either defined by the investigator or by selecting a PheWAS code (phecode) from the catalog (https://phewascatalog.org/phecodes) to use the associated ICD mapping[1 27]. Briefly, the phecode is a published mapping of grouping ICD codes into clinically relevant groups. For example, salmonella pneumonia (PNA) (ICD-9 003.22), PNA due to streptococcus (ICD-9 482.3), etc. are grouped into one phecode “480.1” for “bacterial pneumonia”. Multiple or higher level phecodes can also be used together to represent a broader category, e.g. diabetes mellitus (phecode 250) which includes type 1 diabetes (phecode 250.1) and type 2 diabetes (phecode 250.2). The PheWAS catalog provides a hierarchical roll up information to aggregate broader phenotypes. For a given list of ICD-9 codes selected via phecode mapping or domain expert to represent the phenotype, all codes are aggregated to represent the main ICD feature. Next, the total count of the main ICD feature is extracted from the dataset, denoted by ICD_count_. All ICD-10 codes were first mapped to ICD-9 codes based on the General Equivalence Mapping (GEMS) available at the Centers for Medicare & Medicaid Services[28]. Each ICD-9 code is only counted once per day to reduce the impact of dual coding.

The main NLP feature used in MAP for the phenotype is determined through identifying the medical concepts by mapping relevant clinical terms to the Concept Unique Identifiers (CUIs) listed in the Unified Medical Language System (UMLS). While the main medical concept can be selected by a domain expert, we find it more robust to identify the CUIs through a mapping process utilizing the PheWAS catalogue map, in which a phenotype text string is used to describe each phecode. For each phecode, we identify its corresponding ICD codes along with the phenotype string. We identify a list of CUIs for the phecode through three mapping steps: (i) mapping all relevant ICD-9 codes directly to CUIs, which can be performed directly in UMLS using the (CODE field in the MRCONSO table); (ii) mapping ICD-9 strings to CUIs; (using the STR field in the MRCONSO table) with exact string-matching and (iii) mapping the phenotype string to CUIs (again using the MRCONSO STR field) with exact string-matching (Supplementary Figure 1). Although steps (i) and (ii) typically yield identical lists, (ii) occasionally results in a larger list since an ICD-9 string may be mapped to multiple concepts. Combining all CUIs mapped in the three steps gives a list of CUIs to represent each phecode.

**Figure 1.**
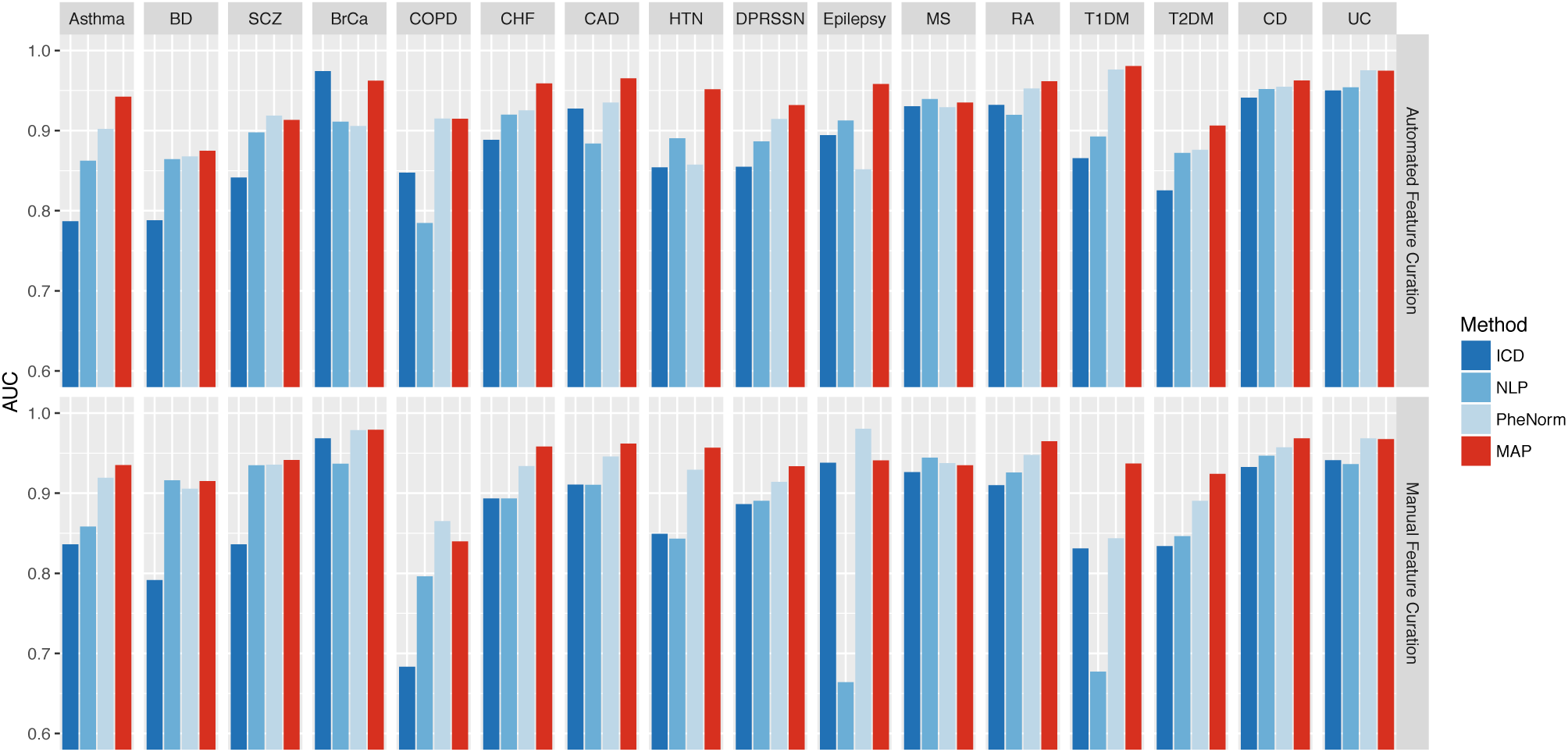
Top panel: Comparison of AUCs with gold standard labels for ICD-9 count, NLP, PheNorm, and MAP for 16 disease phenotypes using the MAP automated feature curation; Bottom panel: Comparison of AUCs for the 16 phenotypes features manual curated by domain experts.

The narrative text notes for all subjects are then processed. Using the CUI list assembled for each phecode, the total number of positive mentions of any clinical terms belonging to the CUIs list, denoted by NLP_count_, is used as the main NLP feature for the phecode. If a phenotype is defined by several phecodes, we take the sum of the corresponding NLP counts as the NLP_count_ feature for the phenotype. The Narrative Information Linear Extraction (NILE)[29] was used to extract concepts in the experiments of this study. Based on previous phenotyping studies, the NLP features extracted were similar regardless of NLP platform used.[10-18]

As demonstrated in the Yu et al.[26] PheNorm study, the level of healthcare utilization contributed noise to algorithm features and dampened their ability to predict the phenotype status. We thus include the total number of narrative notes for each patient, denoted by Note_count_, as a proxy for the healthcare utilization in the MAP algorithm. We also consider modeling the logarithm transformed features, ICD_log_ = log(1 + ICD_count_), NLP_log_ = log(1 + NLP_count_), and Note_log_ = log(1 + Note_count_). Additionally, similar to Yu et al.[26], we combine ICD and NLP counts to create additional features ICDNLP_count_ = 0.5 × (ICD_count_ + NLP_count_) and ICDNLP_log_ = log(1 + ICDNLP_count_).

### Unsupervised MAP Prediction

Using patient level data on the ICD_count_, NLP_count_, ICDNLP_count_, ICD_log_, NLP_log_, ICDNLP_log_, and Note_log_, we separately fitted mixture models to each individual feature along with Note_log_ to obtain the probability that a patient has the phenotype. Specifically, for *X*_count_ ∈ {ICD_count_, NLP_count_, ICDNLP_count_}, we fitted a Poisson mixture model with *X*_count_ | *Y* = *y* ∼ Poisson(*α*Note_log_ + *λ*_*y*_), where *Y* ∈ {0,1} is the unobserved phenotype status. The probability mass function of *X*_count_ is therefore

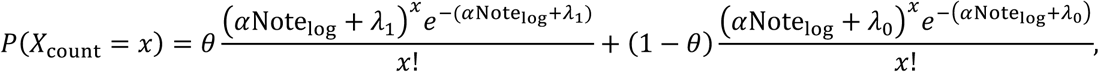

where the parameters *θ*, *α*, *λ*_1_, and *λ*_0_ are estimated with the Expectation-Maximization (EM) algorithm. The parameter *θ* = *P*(*Y* = 1) is the prevalence of the phenotype, *λ*_y_ is essentially the healthcare utilization adjusted mean of *X*_count_ among those with *Y* = *y* for *y* ∈ {0,1}, and *α*Note_log_ is used to mitigate the noise brought into *X*_count_ in predicting the phenotype status, where *α* is generally negative. To increase the robustness of the procedure, we also fit normal mixture models to the log-transformed count data *X*_log_ ∈ {ICD_log_, NLP_log_, ICDNLP_log_} with 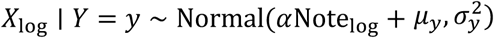 whose probability density function is

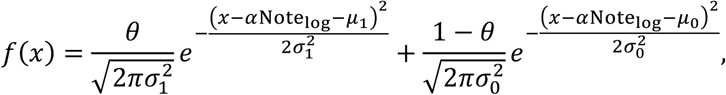

where *θ* = *P*(*Y* = 1), *α*, *μ*_1_, *μ*_0_, 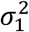, and 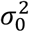 are similarly estimated with the EM algorithm. The parameters *μ*_y_ and 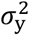 respectively denote the healthcare utilization adjusted mean and variance of *X*_log_ among those with phenotype *Y* = *y*. We fit the above Poisson and log-normal mixture models to the ICD and NLP features as either or both models may provide a good approximation to the observed data and it is unclear which model provides a better approximation for a given dataset, especially in the absence of gold standard labels on *Y*.

To assign a predicted probability to each subject, we calculate the posterior probabilities of having the phenotype given the feature information from each fitted mixture model via the Bayes’ rule. Under each model, the prevalence of the phenotype can be estimated by the average value of the posterior probabilities across all patients. The final MAP algorithm assigns the predicted probability of having the phenotype as the average of the six predicted probabilities. We also estimate the prevalence of the phenotype, *θ*\*, by averaging the prevalence estimates from the six latent mixture model fittings. The prevalence estimate is then used as a threshold to assign a binary classification of whether a subject has the phenotype. Subjects with predicted probabilities above *θ*\* are assigned as having the phenotype with *ŷ* = 1, and those below are assigned as not having the phenotype with *ŷ* = 0 (See Supplementary Method for detailed MAP algorithm description).

### Validation of MAP using real world biorepository data

#### Study Populations

##### Partners Biobank

contains linked EHR and genetic data anchored by two large tertiary care hospitals: Brigham and Women’s Hospital and Massachusetts General Hospital in Boston. A total of 17,805 patients had codified data, NLP data as well as genetic data[30 31]. Additionally, gold standard labels were curated on an average of 545 patients for 16 phenotypes: asthma, bipolar disorder (BD), schizophrenia (SCZ), breast cancer (BrCa), chronic obstructive pulmonary disease (COPD), congestive heart failure (CHF), coronary artery disease (CAD), hypertension (HTN), depression (DPRSSN), epilepsy, multiple sclerosis (MS), rheumatoid arthritis (RA), type I diabetes mellitus (T1DM), type II diabetes mellitus (T2DM), Crohn’s disease (CD) and ulcerative colitis (UC).

##### The Veteran’s Affairs Million Veteran’s Project (VA MVP)

is a longitudinal cohort study, recruiting at approximately 50 VA facilities in the United States[32]. MVP data was used to validate the MAP approach, and to demonstrate the application of MAP to a PheWAS. A total of 330,374 patients had both EHR and genetic data available for the PheWAS analysis.

##### Overview of analyses

The performance of the MAP algorithm was validated in three separate experiments: (i) phenotyping 16 conditions using MAP compared against the gold-standard labels obtained from chart review in the Partners Biobank; (ii) an association study between a low density lipoprotein cholesterol (LDL-C) genetic risk score (GRS) and hyperlipidemia phenotypes; and (iii) a PheWAS in MVP of the interleukin-6R (*IL6R)* genetic variant (Asp358Ala, rs2228145)[33].

##### (i) Testing the performance of MAP against existing approaches for phenotyping across 16 conditions with gold standard labels

In the Partners Biobank, we compared the accuracy of (a) the main ICD feature; (b) the main NLP feature, (c) the PheNorm algorithm, and (d) the MAP algorithm. Similar to previous phenotyping approaches[21 22 26 31], we first applied a filter to create a dataset of subjects who may have the phenotype. This simple filter was ≥1 ICD-9 codes for the phenotype of interest. All subjects who passed this filter consisted of the “filter positive set”. Those without any ICD-9 codes for the phenotype were assigned a probability of 0. Comparisons of the above approaches against gold standard labels were performed among the filter positive set.

For the comparison, the main ICD features using in MAP were defined in two ways, either a list of ICD-9 codes defined by domain experts or generated using the semi-automated process as part of MAP. Similarly, the main NLP feature was defined either as a list created by domain experts or using the mapping steps described above. The optional denoising step in the PheNorm algorithm was not applied because it requires additional candidate features beyond the main ICD and NLP features, limiting its ability to scale up for application to the PheWAS.

We evaluated the overall performance of the main ICD, main NLP, PheNorm, and MAP against the gold standard labels for the 16 phenotypes using the area under the receiver operating characteristic curve (AUC). In addition, we compared the MAP classification rule which automatically provides an estimated threshold to classify subjects in phenotype yes/no categories. The performance of the MAP-based binary yes/no classification were compared to phenotypes defined using ≥2 ICD-9 codes. We reported the positive predictive value (PPV) or precision, negative predictive value (NPV), and F-score, against the 16 phenotypes with gold standard labels.

##### (ii) MAP for genetic association studies

Using genetics and EHR data from Partners Biobank, we performed a genetic association study between a previously published LDL-C GRS[34 35] and the disease phenotype of hyperlipidemia defined either using MAP predicted probabilities or based on having ≥2 ICD9 codes for hyperlipidemia (ICD-9 272.x), corresponding to the phecode 272.1[1 36]. The association analyses were performed by fitting a logistic regression model regressing the MAP probabilities or ICD-9 based binary status for hyperlipidemia against the LDL GRS, adjusted for age, gender, and self-reported race.

##### (iii) MAP for PheWAS

In MVP, we performed a PheWAS for the IL6R variant, rs2228145. Individuals with the IL6R variant have profiles similar to individuals on IL6R antagonists, tocilizumab and sarilumab; both have been indicated for the treatment of rheumatoid arthritis (RA), and tocilizumab for giant cell arteritis (GCA). A recently published standard PheWAS based on ICD codes using MVP data found 22 phenotypes significantly associated with IL6R at the false discovery rate of 0.05[33]. While this SNP was found to be associated with vascular and cardiac diseases, its associations with RA and GCA were not statistically significant. In this analysis, we tested whether MAP compared to the standard PheWAS method improved the power to detect these expected associations.

For each phecode, we predicted the presence of the phenotype based on either the MAP algorithm or the ICD-9 code. Since MAP provided predicted probabilities of having the phenotype, we performed PheWAS directly relating the predicted probabilities from MAP to the genetic variant by fitting a quasi-binomial model. For both PheWAS methods, we adjusted for age, gender, and self-reported race; the robust sandwich estimator[37] for the variance. We performed PheWAS on 1606 phenotypes, defined as the phecodes with prevalence >0.1% in MVP.

## RESULTS

### The performance of MAP against existing approaches for phenotyping across 16 phenotypes

The ICD-9 and NLP concepts selected using the proposed semi-automated method as part of MAP yielded phenotype algorithms with similar AUCs to those developed using features selected by domain experts (Figure 1). The average AUC of the 16 MAP algorithms developed using ICD-9 + NLP features extracted using the MAP automated feature curation was 0.943, compared to 0.941 for algorithms developed from manual feature curation (Supplementary Table 1). The average AUCs of the individual ICD and NLP features created by the semi-automated process were 0.881 and 0.896, slightly higher than that of those curated manually, 0.873 and 0.870, respectively. We focused on discussing the results using ICD-9 and NLP concepts selected by the proposed semi-automated method in the following sections.

Across the 16 phenotypes with gold standard labels, MAP classified the phenotypes with higher accuracy than ICD-9 or NLP in 13 out of the 16 cases (Figure 1, top panel). Also, the MAP algorithm outperformed PheNorm algorithm in most phenotypes and the most significant improvement in AUC were observed in Epilepsy and HTN (Supplementary Table 1). Compared to the ICD-9 feature, NLP feature, and PheNorm, the MAP algorithm improved the AUC significantly by about 0.062 (p-value = 2.37 × 10^−9^), 0.047 (p-value = 2.91× 10^−11^), and 0.027 (p-value = 7.21 × 10^−7^) on average, respectively (Supplementary Table 1).

The MAP-based binary yes/no classification also had higher accuracy than phenotypes defined by using ≥2 ICD-9 codes when evaluated against a gold standard. Across the 16 phenotypes, MAP classified all phenotypes with a higher PPV (precision) than ICD-9 codes while maintaining a comparable NPV (Figure 2). For instance, the PPVs for the MAP and ICD-9 code were 0.947 versus 0.621 for RA, 0.876 versus 0.569 for CAD (Supplementary Table 2). Accounting for the trade-off between sensitivity (recall) and PPV (precision), the MAP classification attained a higher F-score than the simple ICD-9 rule across most phenotypes, resulting in an average difference in F-score of 0.076 (p-value = 4.11 × 10^−9^). In addition, MAP had larger AUCs (AUC here is for binary classifier) than ICD-9 codes across 16 phenotypes with average difference of 0.136 (p-value = 4.47 × 10^−23^). Similarly, MAP had better performance than binary classifier defined by NLP features (Supplementary Table 2).

**Figure 2.**
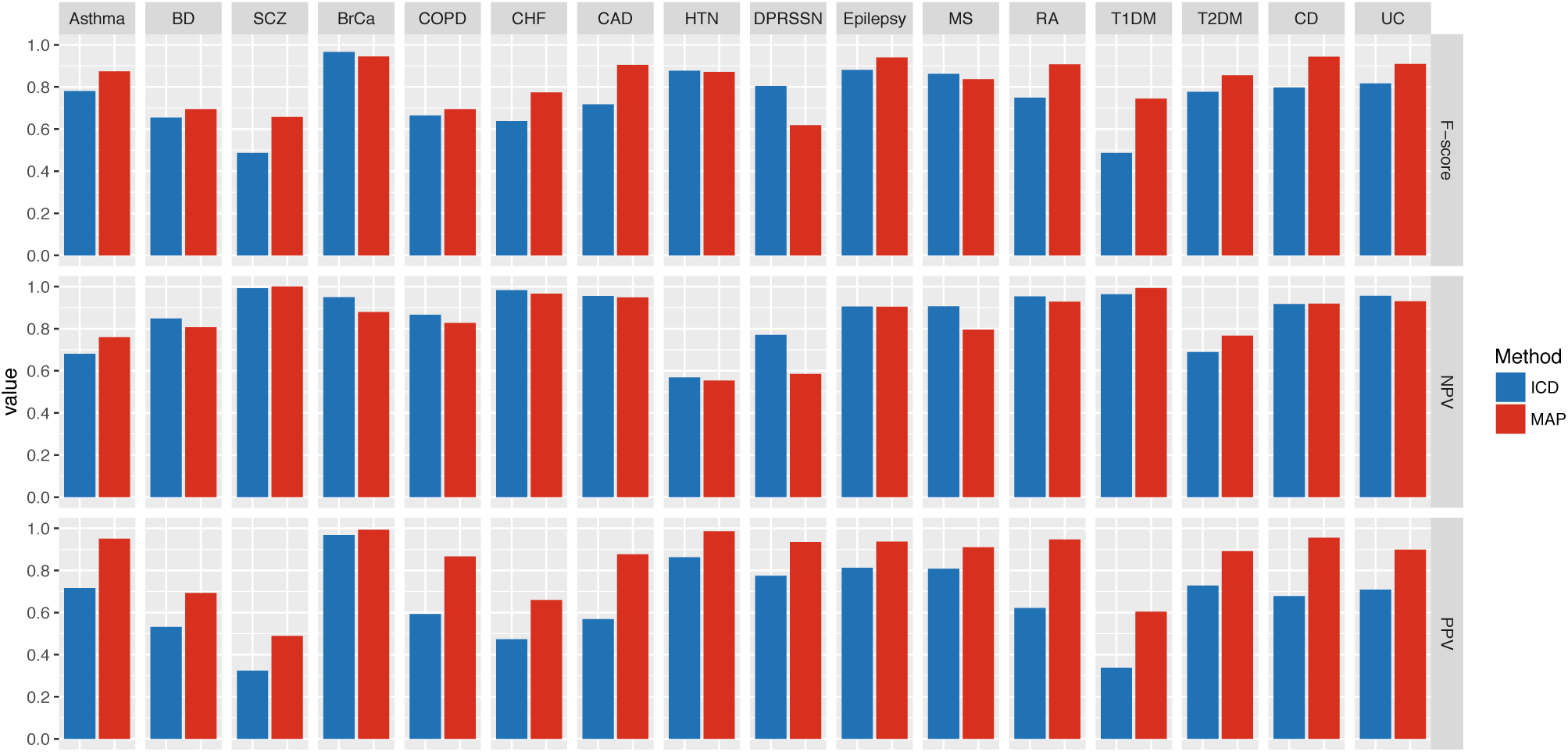
Performance of phenotype classification using MAP compared to ICD-9 codes for 16 phenotypes with gold-standard labels. [F-score, negative predictive value (NPV), positive predictive value (PPV, precision)].

### Genetic Association Between LDL GRS and Hyperlipidemia in the Partners Biobank

Among Partners Biobank subjects, a total of 8,226 (46.2%) out of 17,805 subjects had ≥2 ICD-9 codes for hyperlipidemia. The estimated odds ratio (OR) between the LDL-C GRS and hyperlipidemia was 1.007 (p-value = 0.015) when hyperlipidemia was defined using the standard ICD-9 code thresholding approach. In contrast, the estimated OR was 1.010 (p-value = 0.0001) when the hyperlipidemia phenotype was defined using MAP, suggesting that MAP had improved power for association test compared to ICD-9 code.

### PheWAS using VA MVP Data

In MVP, we performed both MAP based and ICD-9 based PheWAS of *IL6R* rs2228145. The minor allele frequency for *IL6R* rs2228145 (risk allele: A; Asp358Ala) was 0.35. The PheWAS results based on the MVP data are shown in Figure 3. Detailed results for the association pairs that attained significant p-values after Bonferroni correction based on either MAP or ≥2 ICD-9 codes are shown in Supplementary Table 3. Compared to the standard ICD-9 code based PheWAS, MAP yields a similar number of significant associations when using Bonferroni correction, 13 for MAP and 12 for ICD. MAP, using the predicted probability as the outcome, generally detected stronger associations with odds ratios in larger magnitude and smaller p-values when compared to the results based on the standard PheWAS. For example, the estimated OR between IL6R SNP and ischemic heart disease was 0.945 (p-value = 3.89 × 10^−19^) for MAP and 0.955 (p-value = 3.59 × 10^−12^) for ICD. Among the 17 phenotypes that were significantly associated with IL6R by either MAP or standard PheWAS, the average negative log10 p-value was 7.40 for MAP and 6.78 for ICD while the average absolute log odds ratio was estimated as 0.067 by MAP but 0.064 by ICD.

**Figure 3.**
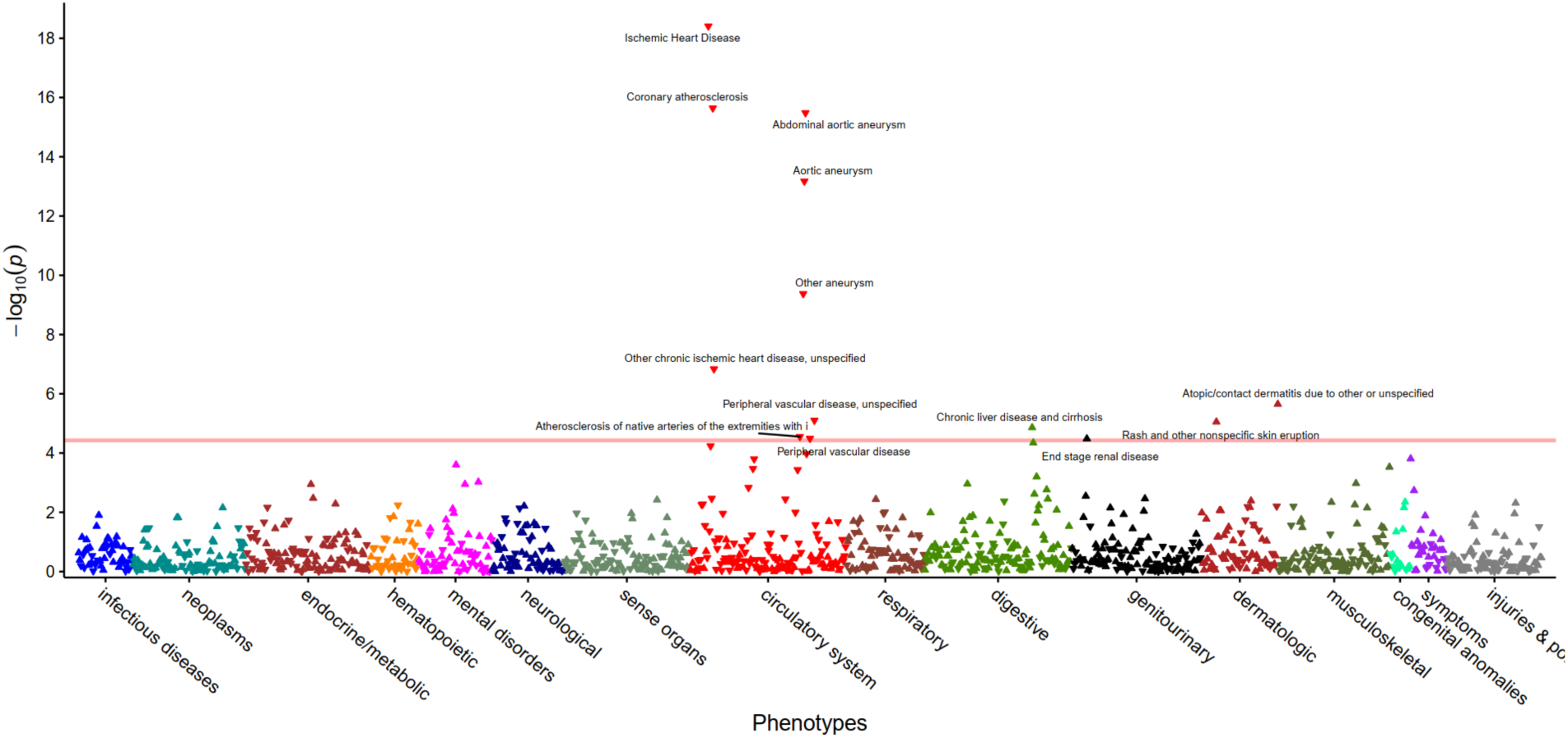
PheWAS results using MAP defined phenotypes for the IL6R SNP. Phenotypes significantly associated with IL6R after Bonferroni correction are annotated.

The IL6R antagonists were approved for treating rheumatoid arthritis (RA), giant cell arteritis (GCA), and systemic juvenile idiopathic arthritis (sJIA), also known as juvenile rheumatoid arthritis (JRA). However, the associations between *IL6R* and these three phenotypes were not detected in the VA MVP PheWAS due to the low prevalence and low accuracy of the ICD codes^[33]^. The *IL6R* PheWAS in MVP using ICD codes to define phenotypes found the following associations: GCA 0.926 (p-value = 0.279), RA 0.984 (p-value = 0.402), and juvenile RA (JRA) 1.082 (p-value = 0.684). Using MAP based phenotype definitions, the following associations were observed with IL6R: GCA 0.883 (p-value = 0.082), RA OR 0.986 (p-value = 0.494) and JRA OR 1.152 (p-value = 0.509) (Supplementary Table 4). Defining phenotypes with MAP showed a stronger association between *IL6R* and reduced risk of GCA whereby the magnitude of the reduced risk was higher with MAP as well as the level of significance. For JRA and RA, both MAP and ICD did not detect the associations, likely due to the inaccuracy in the coding and specific characteristics of the VA population.

We also validated the association of GCA, RA and juvenile RA (JRA) using the Partner’s biobank. Using an ICD based definition for phenotypes the associations with IL6R showed the following: GCA OR 0.870 (p-value = 0.320), RA OR 0.897 (p-value = 0.008), and JRA OR 0.745 (p-value = 0.050). Using MAP to define the phenotypes, the estimated OR was 0.853 (p-value = 0.269) for GCA, 0.888 (p-value = 0.007) for RA, and 0.721 (p-value = 0.041) for JRA (Supplementary Table 4). These results again suggest improved power in detecting associations using the MAP approach.

## DISCUSSION

In summary, MAP provided three major advancements for high-throughput phenotyping. First, MAP provides a systematic automated approach for integrating narrative information using NLP with ICD codes for use in a high-throughput phenotype algorithm pipeline. Second, MAP automates selection of the NLP concepts for a given phenotype, traditionally a labor-intensive manual process. Third, built into MAP is the ability to assign a threshold to provide a binary yes/no classification for each phenotype. This was accomplished using the existing EHR data, thereby bypassing another traditionally rate limiting step, medical record review to determine the threshold probability above which a subject is considered to have the phenotype. This high degree of automation allows the integration of more data extracted using NLP into the algorithms, resulting in improvements in accuracy, while maintaining the scalability enabling large scale phenotype screens such as the PheWAS. The improved power and scalability will also enable more efficient and less biased epidemiological studies since MAP can be used to accurately annotate both the main phenotype of interest and other risk factors or covariates. In addition, the scalability of MAP is enhanced by its efficiency in computation time. For MVP cohort, it took about 1 minute per phenotype to run MAP utilizing one core on an Intel Xeon 5650 6-Core CPU (2.66 GHz) and it took NILE about 1 second to process 1000 notes that have at least 500 characters using the dictionary of 43,291 terms and 12,621 CUIs.

Similar to an existing high-throughput phenotyping approach such as PheNorm[26] and others[24]. MAP can rank order subjects according to the likelihood of having the phenotype. However, MAP also provides a predicted probability of having the phenotype as well as an automated method to a determine a threshold for each phenotype, facilitating classification into the binary phenotype yes/no categories. This ability to define a phenotype in two ways provides options for investigators depending on the study question. Using a probability of phenotype rather than a binary yes/no classification allows investigators to consider a phenotype as a continuous trait. This characteristic improved power to detect associations[38] compared to the standard PheWAS approach, which uses a binary trait, with higher effect sizes and smaller p-values. From our study using Partners Biobank, using MAP defined hyperlipidemia yielded much more significant result than that defined by ICD code, regarding the association between the LDL GRS and hyperlipidemia. This further confirms that using MAP defined phenotypes can improve the statistical power of downstream association studies as well as more accurate estimate of the effect sizes compared to ICD based analyses.

The option of classifying subjects into yes/no binary phenotype categories is also a step not available in PheNorm or other existing unsupervised algorithms. The ability to generate a data driven threshold that is adaptive to the specific phenotype and the healthcare system is also a desirable feature. Existing methods largely use a common threshold (e.g. 1 or 2 for ICD codes) but may result in poor performance for specific phenotypes or poor portability due to heterogeneity in healthcare systems[10]. Prevalence estimates using MAP, ICD codes, and chart review are provided in Supplementary Table 2, which suggests that the MAP estimates are more consistent with that based on chart review. Throughout our experiments, we assign a probability of 0 for patients in the filter negative set and evaluation on the filter positive set. The filter defined as having at least 1 ICD code generally has a high NPV. For example, in Partner’s Biobank, the NPV for having no ICD code attained an average of 99% across the 16 phenotypes with 10 of which reaching 100%. In addition, the average NPV across 16 phenotypes for all patients (including both filter negative and filter positive patients) was 0.994 for both MAP and ICD-9 code (Supplementary Table 5). These suggest that assigning a probability of 0 for filter negative patients is safe and reasonable. However, alternative filters may have better performance and our MAP approach is not restricted to any specific filter. For under coded diseases phenotypes such as suicide ideation, filters based on both NLP and ICD may be attain higher accuracy.

A key part of MAP is the use of automated steps to identify the relevant ICD or NLP features for any given phenotype algorithm. The algorithms developed using ICD and NLP features selected using the MAP process had comparable or slightly better AUCs than manually curated features selected by domain experts across the 16 phenotypes. Additionally, the experiment shows that MAP was more robust than PheNorm. MAP uses model averaging over three types of features (ICD, NLP, and ICD+NLP) and two types of distributions (Poisson and log-normal) for algorithm development; PheNorm uses only the normal distribution and relies on majority voting. The additional models used by MAP improves the robustness of the algorithm. Specifically, to see the impacts of each single model and healthcare utilization on the performance of the MAP algorithm, we compared the performance of our proposed MAP algorithm to several simplified versions of MAP algorithm by removing some of the components for phenotyping the 16 diseases. From the results (Supplementary Tables 1 and 2), we found that the performance of MAP algorithm decreased when some of the components were excluded. To aggregate the results from all six models, MAP currently uses a simple model average of the predicted probabilities, in part due to the difficulty in ascertaining the performance of mixture models in the absence of labels. Data adaptive weighting can potentially be used to further improve the MAP performance.

It is worth noting that the denoising step of the original PheNorm was not used in this study. The denoising step can potentially improve the accuracy of PheNorm when the main ICD and NLP features are not sufficiently predictive, for example in the cases of asthma with automated features and T1DM with manual features. In comparison, MAP had a more robust performance and can attain reasonable accuracy in the absence of additional features and is thus highly scalable. However, it is plausible that incorporating additional features may further improve the accuracy of MAP for certain phenotypes, which warrants further research.

Compared to ICD based analyses, the improved accuracy of MAP translated into improved power in both association studies and a PheWAS using real world cohort data. From the genetic association study example between the LDL GRS and hyperlipidemia in the Partners Biobank (n=17,805), we observed that improving the accuracy of phenotype definitions is particularly important when the sample size is relatively small. The association between LDL GRS and hyperlipidemia was much more significant when the phenotype is defined by MAP compared to that defined by ICD-9 codes alone. In MVP and Partners Biobank, MAP provided improved power to detect associations between *IL6R* and uncommon phenotypes where the accuracy of the ICD code was relatively low.

## CONCLUSION

Using a relatively large number of phenotypes with gold standard labels for validation, we demonstrated that the proposed MAP algorithm achieves more efficient, robust, and accurate phenotyping compared to existing approaches. One distinct advantage to MAP is the process of providing automated selection of both ICD and NLP features and integration of multiple feature types and distributions. We validated the MAP approach in two independent biobanks, demonstrating that MAP annotated phenotypes are comparable to those developed using manual approaches, can replicate known associations, and improve power in smaller datasets. Finally, MAP provides two forms of phenotype data for studies, in a traditional phenotype yes/no format or as a probability of a phenotype. The MAP high-throughput phenotyping approach integrating ICD-9 and key NLP concepts has direct applications in large scale association studies such as PheWAS in biorepositories linked with EHR data. This promising approach can also improve the resolution, particularly among less common or poorly coded conditions, to study the relationships across multiple phenotypes in one study.

## Funding

This work was supported in part by the U.S. National Institutes of Health Grants P30-AR072577, U54-HG007963, U01-HG008685 and RO1-HG009174, National Natural Science Foundation of China Grant 11801301, and National Key R&D Program of China Grant 2018YFC0910404.

## Competing Interests

None.

## Contributorship

All authors made substantial contributions to: conception and design; acquisition, analysis and interpretation of data; drafting the article or revising it critically for important intellectual content; and final approval of the version to be published.

## Notes

\# Part of this research is based on data from the Million Veteran Program, Office of Research and Development, Veterans Health Administration, and was supported by award #MVP000. This publication does not represent the views of the Department of Veterans Affairs or the United States Government.

