## Supplementary figure for "High-throughput Multimodal Automated Phenotyping (MAP) with Application to PheWAS"

Supplementary Figure 1. Illustration of CUI mapping for a given phecode using its associated ICD9 codes, ICD9 strings and PheWAS strings: (a) flow chart for the mapping process; and (b) illustration of CUI mapping process for phecode “174”.
