## Supplementary method for "High-throughput Multimodal Automated Phenotyping (MAP) with Application to PheWAS"

### Detailed MAP Algorithm Description:

1. Given the input variables $\{\text{ICD}_{\text{count}}, \text{NLP}_{\text{count}}, \text{ICDNLP}_{\text{count}}, \text{Note}_{\text{log}}\}$, a larger set of variables are created, i.e. $\{\text{ICD}_{\text{count}}$, $\text{NLP}_{\text{count}}$, $\text{ICDNLP}_{\text{count}}$, $\text{ICD}_{\text{log}}$, $\text{NLP}_{\text{log}}$, $\text{ICDNLP}_{\text{log}}$, $\text{Note}_{\text{log}}\}$.

2. Create filter positive cohort based on the input variables. For example, the filter $\text{ICD}_{\text{count}} \geq 1$ can be used to define filter positive patients, as we did in the paper.

3. For patients in the filter positive cohort:

   (a) For each $X_{\text{count}} \in \{\text{ICD}_{\text{count}}, \text{NLP}_{\text{count}}, \text{ICDNLP}_{\text{count}}\}$, fit a Poisson mixture model adjusting for $\text{Note}_{\text{log}}$ and then obtain the corresponding posterior probability of each patient having the disease.

   (b) For each $X_{\text{log}} \in \{\text{ICD}_{\text{log}}, \text{NLP}_{\text{log}}, \text{ICDNLP}_{\text{log}}\}$, fit a Normal mixture model adjusting for $\text{Note}_{\text{log}}$ and then obtain the corresponding posterior probability of each patient having the disease.

   (c) Let $\widehat{p}_{im}$ $(i = 1, ..., N, m = 1, ..., 6)$ be the posterior probability of patient $i$ obtained from the $m_{\text{th}}$ model fitting and $\widehat{q}_{im} = \Phi^{-1}(\widehat{p}_{im})$ be the transformed variable, where $\Phi(\cdot)$ is the cumulative distribution function of a standard Normal distribution and $N$ is the number of filter positive patients.

   (d) Derive the pre-score $\widetilde{p}_i = \frac{1}{2}\left(\frac{1}{6}\sum_{m=1}^{6}\widehat{p}_{im} + \Phi(\frac{1}{6}\sum_{m=1}^{6}\widehat{q}_{im})\right)$ for each patient and calculate an initial estimate of prevalence as $\widehat{\theta}_1 = \frac{1}{N}\sum_{i=1}^{N}\widetilde{p}_i$.

   (e) To get a robust estimate of the prevalence, we use K-means clustering to group patients into two clusters based on the two variables, that is the average count of $\text{ICD}_{\text{count}}$, $\text{NLP}_{\text{count}}$, and $\text{ICDNLP}_{\text{count}}$ and the average of the transformed variables $\frac{1}{3}\sum_{m=4}^{6}\widehat{q}_{im}$ (models 4, 5, and 6 are the Normal mixture models). Another estimate of the prevalence $\widehat{\theta}_2$ is then obtained as the proportion of patients in the cluster having larger average count. Final estimate of prevalence is then calculated as $\widehat{\theta} = \frac{1}{2}(\widehat{\theta}_1 + \widehat{\theta}_2)$.

   (f) The final probability of each patient having the disease $\widehat{P}_i = g^{-1}(g(\widetilde{p}_i) - c)$, where $g(\cdot)$ is the logit function defined as $g(p) = \log(\frac{p}{1-p})$ and $c$ is calculated by solving the equation $\frac{1}{N}\sum_{i=1}^{N} g^{-1}(g(\widetilde{p}_i) - c) = \widehat{\theta}$. This step is essentially to re-scale the predicted probabilities so that its mean matches the prevalence estimate.

4. For patients in the filter negative cohort, we set $\widehat{P}_i = 0$.
